# Estimated connectivity networks outperform observed connectivity networks when classifying people with multiple sclerosis into disability groups

**DOI:** 10.1101/2021.06.07.447376

**Authors:** Ceren Tozlu, Keith Jamison, Zijin Gu, Susan A. Gauthier, Amy Kuceyeski

## Abstract

**Background:** Multiple Sclerosis (MS), a neurodegenerative and neuroinflammatory disease, causing lesions that disrupt the brain’s anatomical and physiological connectivity networks, resulting in cognitive, visual and/or motor disabilities. Advanced imaging techniques like diffusion and functional MRI allow measurement of the brain’s structural connectivity (SC) and functional connectivity (FC) networks, and can enable a better understanding of how their disruptions cause disability in people with MS (pwMS). However, advanced MRI techniques are used mainly for research purposes as they are expensive, time-consuming and require high-level expertise to acquire and process. As an alternative, the Network Modification (NeMo) Tool can be used to estimate SC and FC using lesion masks derived from pwMS and a reference set of controls’ connectivity networks.

**Objective:** Here, we test the hypothesis that estimated SC and FC (eSC and eFC) from the NeMo Tool, based only on an individual’s lesion masks, can be used to classify pwMS into disability categories just as well as SC and FC extracted from advanced MRI directly in pwMS. We also aim to find the connections most important for differentiating between no disability vs evidence of disability groups.

**Materials and Methods:** One hundred pwMS (age:45.5 *±* 11.4 years, 66% female, disease duration: 12.97 ± 8.07 years) were included in this study. Expanded Disability Status Scale (EDSS) was used to assess disability, 67 pwMS had no disability (EDSS *<* 2). Observed SC and FC were extracted from diffusion and functional MRI directly in pwMS, respectively. The NeMo Tool was used to estimate the remaining structural connectome (eSC), by removing streamlines in a reference set of tractograms that intersected the lesion mask. The NeMo Tool’s eSC was used then as input to a deep neural network to estimate the corresponding FC (eFC). Logistic regression with ridge regularization was used to classify pwMS into disability categories (no disability vs evidence of disability), based on demographics/clinical information (sex, age, race, disease duration, clinical phenotype, and spinal lesion burden) and either pairwise entries or regional summaries from one of the following matrices: SC, FC, eSC, and eFC. The area under the ROC curve (AUC) was used to assess the classification performance. Both univariate statistics and parameter coefficients from the classification models were used to identify features important to differentiating between the groups.

**Results:** The regional eSC and eFC models outperformed their observed FC and SC counterparts (p-value*<*0.05), while the pairwise eSC and SC performed similarly (p=0.10). Regional eSC and eFC models had higher AUC (0.66-0.68) than the pairwise models (0.60-0.65), with regional eFC having highest classification accuracy across all models. Ridge regression coefficients for the regional eFC and regional observed FC models were significantly correlated (Pearson’s *r* = 0.52, p-value *<* 10e-7). Decreased estimated SC node strength in default mode and ventral attention networks and increased eFC node strength in visual networks was associated with evidence of disability.

**Discussion:** Here, for the first time, we use clinically-acquired lesion masks to estimate both structural and functional connectomes in patient populations to better understand brain lesion-dysfunction mapping in pwMS. Models based on the NeMo Tool’s estimates of SC and FC better classified pwMS by disability level than SC and FC observed directly in the individual using advanced MRI. This work provides a viable alternative to performing high-cost, advanced MRI in patient populations, bringing the connectome one step closer to the clinic.

**Highlights:**

- We compared the accuracy of models based on observed functional connectivity (FC) and structural connectivity (SC) networks extracted from advanced MRI and estimated FC and SC networks derived using only lesion masks from conventional MRI in classifying people with multiple sclerosis (pwMS) into disability groups.
- Estimated SC and FC generally outperformed observed SC and FC in classifying pwMS into no disability vs evidence of disability groups, with regional estimated SC and FC having the best performance.
- Increased estimated FC node strength of regions in the visual network was associated with disability.
- Decreased estimated SC node strength of regions in the default mode and ventral attention networks was associated with disability.
- Despite their varied sources of origin, feature weights for the regional estimated FC and the regional observed FC classification models was significantly correlated (Pearson’s *r* = 0.52, p-value *<* 10e-7).

## 1 Introduction

Multiple sclerosis (MS) is a disease that is largely characterized by brain lesions, detected using clinical MRI, that cause cognitive impairment and/or physical disability. The number and size of lesions, however, is not always proportional to an individual’s disability (1); recent work has demonstrated that lesions’ disruption to structural and physiological connectivity networks may be more important for accurate lesion-dysfunction mapping (2; 3). Connectomics, or brain connectivity network analysis, provides a promising tool to directly measure the effect of MS-related pathology and to capture the reorganization mechanisms in response to pathology. Having a more accurate lesion-dysfunction map, particularly a lesion-future dysfunction map, would allow development of more accurate prognoses and could be used to personalize treatment plans to improve clinical care of people with MS (pwMS).

Advanced neuroimaging techniques such as diffusion magnetic resonance imaging (dMRI) and functional MRI (fMRI) are commonly used to quantify the structural (white matter) connectome (SC) and the functional (co-activation) connectome (FC). Previous studies have shown that SC metrics can successfully differentiate healthy controls (HC) from pwMS, classify pwMS who showed different clinical phenotypes, and be used to assess its relationship with disability (4; 5; 6). However, tractography can be challenging in areas of complex fiber orientations and can produce false-positive and -negative connections. Lesions, pathologies in normal-appearing white matter, inflammation, demyelination and axonal loss may decrease SNR in dMRI and make existing tractography issues even more pervasive in pwMS (7; 8; 9). In terms of measuring FC in pwMS, inflammation induces alterations in synaptic transmission (10; 11) and thus may affect the BOLD signal in non-trivial ways (12; 13). FC changes in pwMS relative to controls may also be driven by compensation (14), maladaptation (15) or both (16), making the mapping of FC to dysfunction even more complicated. Therefore, observed SC and FC extracted from advanced MRI in pwMS may not be accurately representing underlying brain networks and/or be informative of disability.

In addition to the above outlined challenges using advanced MRI in pwMS, the imaging techniques themselves are expensive, time-consuming and require a high level of expertise. Therefore, alternative techniques that use only conventional, clinically-acquired MRI may allow more widespread use of SC and FC measures for understanding the mechanisms and connectome-behavior mapping in pwMS (17; 18). As an alternative to performing tractography in pwMS, lesion masks extracted from conventional MRI can be used to calculate estimated structural connectivity (eSC) via the Network Modification (NeMo) Tool (19). The recently updated NeMo Tool version 2.0 (https://kuceyeski-wcm-web.s3.useast-1.amazonaws.com/upload.html) uses tractography results from 420 unrelated Human Connectome Project controls as a reference; a lesion mask from a pwMS is used to remove “damaged” streamlines that pass through the lesions from the reference tractographies and the remaining streamlines are used to obtain eSC. A recent work introduced a deep neural network trained to predict FC from SC (20). Using their framework, we created a neural network trained on the NeMo Tool’s 420 control reference subjects to predict FC from SC. This neural network was then integrated into the NeMo Tool so that it could take its eSC and produce the corresponding estimated FC (eFC), a functionality which as far as we know is not available in other tools that estimate structural disconnectivity (17; 18). To summarize, the NeMo Tool version 2.0 uses only an individual’s lesion mask to produce estimates of their SC and FC networks.

Observed connectivity from advanced imaging and estimated disruptions in connectivity based on conventional imaging have been independently associated with disability and cognitive impairment in pwMS (3; 21; 22; 14). A recent study (23) showed structural disconnectivity due to stroke lesions is a better predictor than indirect functional disconnectivity, estimated using symptom-lesion network mapping (24), in predicting multiple functional deficits in stroke. Moreover, the same study showed observed FC from fMRI was superior than indirect functional disconnectivity in the subset of the stroke patients. However, no study has yet compared the ability of observed and estimated SC/FC metrics in classifying pwMS into disability categories.

In this paper, models based on eSC and eFC derived from clinical MRI-derived lesion masks and the NeMo Tool were compared with models based on SC and FC extracted directly from advanced MRI in their ability to classify pwMS into disability groups. Specifically, the classification performance of models based on pairwise and regional SC, FC, eSC, and eFC metrics were compared. We hypothesized that models based on eSC and eFC metrics would perform at least as well as the SC and FC metrics from advanced imaging. Second, the most important pairwise and region-wise connections in the classification task were quantified.

## 2 Materials and Methods

### 2.1 Subjects

One-hundred pwMS (age:45.5 *±* 11.4, 66% females) with a diagnosis of clinically isolated syndrome (CIS) or MS (7 CIS, 88 relapsing-remitting, 5 primary or secondary progressive MS) were enrolled in our study. Possible participants were excluded if they had contraindications to MRI. Demographic data was collected (age, sex, race, clinical phenotype, and disease duration) and subjects underwent an MRI scan. Extended Disability Status Score (EDSS) was used to quantify disability, where an EDSS *<* 2 was considered no disability and EDSS *≥* 2 considered evidence of disability. This threshold was defined based on the fact that pwMS having EDSS values of 0-1.5 present some abnormal signs/symptoms but do not have functional disability. The spinal cord lesion number was included in the patients clinical radiology report and this was used to estimate spinal cord burden in patients. All studies were approved by an ethical standards committee on human experimentation, and written informed consent was obtained from all patients.

### 2.2 Image acquisition, processing and connectome extraction

MRI data were acquired on a 3T Siemens Skyra scanner (Siemens, Erlangen, Germany) with a 20-channel head-neck coil and a 32-channel spine-array coil. Anatomical MRI (T1/T2/T2-FLAIR, 1 mm^3^ iso-voxel), resting-state fMRI (6 min, TR = 2.3 s, 3.75 x 3.75 x 4 mm voxels) and diffusion MRI (55 directions HARDI, b=800, 1.8 x 1.8 x 2.5 mm voxels) acquisitions were performed. Sagittal STIR images were acquired for identification of spinal lesions (TR = 3.5 s, TI=220ms, TE=45ms, in-plane resolution 0.43mm, FOV=22mm, slice thickness 3mm). Multi-echo 2D GRE field maps were collected for use with both fMRI and diffusion MRI (0.75 x 0.75 x 2 mm voxels, TE1=6.69 ms, *l:;*TE=4.06 ms, number of TEs = 6).

#### 2.2.1 Structural and functional connectivity extraction

White matter (WM) and gray matter (GM) were segmented and GM further parcellated into 86 regions of interest (68 cortical and 18 subcortical/cerebellar) using FreeSurfer (25). As described elsewhere (26), fMRI preprocessing included simultaneous nuisance regression and removal of WM and cerebrospinal fluid (CSF) effects (27), followed by band-pass filtering (0.008-0.09Hz) using the CONN v18b toolbox (28) and SPM12 in Matlab. Nuisance regressors included 24 motion parameters (6 rotation and translation, temporal derivatives, and squared version of each) and the top 5 eigenvectors from eroded masks of both WM and CSF. The mean fMRI signal over all voxels in a region was calculated and the mean regional time series correlated (Pearson’s correlation) between every pair of regions to obtain pairwise FC matrices. Regional FC node strengths were calculated by taking the sum of the columns in the FC matrix after removing the negative entries.

Diffusion MRI was interpolated to isotropic 1.8mm voxels, and then corrected for eddy current, motion and EPI-distortion with the eddy command from FSL 5.0.11 (29) using the outlier detection and replacement option (30). MRtrix3Tissue (https://3Tissue.github.io), a fork of MRtrix3 (31) was used to estimate a voxel-wise single-shell, 3-tissue constrained spherical deconvolution model (SS3T-CSD) and then compute whole-brain tractography for each subject. We performed deterministic (sd-stream) tractography with MRtrix3 (32) (33), after which the SIFT2 global filtering algorithm (34) was applied to account for bias that exists in greedy, locally-optimal streamline reconstruction algorithms. The SC matrix was constructed by taking the sum of the SIFT2 weights of streamlines connecting pairs of regions and then dividing by the sum of the two regions’ volumes. In addition to the pairwise SC measures, regional SC node strength was quantified by taking the sum of each of the columns in the SC matrix.

#### 2.2.2 Lesion mask creation

The WM hyperintensity lesion masks were created by running the T2 FLAIR images through the Lesion Segmentation Tool (LST) and were further hand-edited as needed. T2 FLAIR-based lesion masks were transformed to the individual’s T1 native space using the inverse of the T1 to GRE transform and nearest-neighbor interpolation. Individual T1 images were then normalized to MNI space using FSL’s linear (FLIRT) and non-linear (FNIRT) transformation tools (http://www.fmrib.ox.ac.uk/fsl/index.html); transformations with nearest-neighbor interpolation were then applied to transform the native anatomical space T2FLAIR lesion masks to MNI space. The transformations were concatenated (T2FLAIR to T1 to MNI) to minimize interpolation effects. Lesions were manually inspected after the transformation to MNI space to verify the accuracy of coregistration.

#### 2.2.3 Network Modification Tool

The NeMo Tool (19) used each subject’s MNI-space T2 FLAIR lesion mask to calculate eSC and eFC using a reference database of SCs and FCs from 420 unrelated controls from the Human Connectome Project (see supplementary material for details). The NeMo Tool’s original functionality was to estimate SC (or structural disconnectivity) due to a lesion mask by identifying streamlines from each healthy control tractography set that pass through the lesion mask, and then recording the gray matter regions they connect. The eSC matrix is then the sum of the SIFT2 weights of the remaining streamlines connecting pairs of regions, divided by the sum of the two regions’ volumes averaged over the 420 individuals in the reference set. Regional eSC node degree was taken to be the sum of the columns in the eSC matrix.

In addition, we extended the NeMo Tool’s functionality to create eFC from lesion masks by first creating a deep neural network that can predict FC from SC using a previously published approach (20). We used the NeMo Tool’s 420 subjects to train, validate and test the model, which was optimized to minimize the difference between predicted and observed FC but also minimize the l1 distance between inter-subject similarity of the predicted FC and observed FC across the individuals in the training set. The model, which takes in the upper triangular part of the SC matrix, is a feed-forward fully connected neural network comprised 8 hidden layers with 1024 neurons and a dropout rate of 0.5 for each layer. We used leaky rectified linear unit (ReLU, leakage parameter set to 0.2) and hyperbolic tangent (tanh) as the activation functions and initialized the weights using the Xavier algorithm. Adam was used as our optimizer. The hyperparameters in the loss function include *λ*, the regularization parameter, and *γ*, the parameter that modeled averaged the inter-subject correlation of the observed FC of each individual in the training set, were determined by grid search in the interval of [0.1, 0.7] with steps of 0.1 for *γ*, and across values of [0.0001, 0.0005, 0.001, 0.005, 0.01, 0.05, 0.1, 0.5, 1] for *λ*. The best value for these parameters was found to be *γ* = 0.2 and *λ* = 0.01. We trained the model using 340 individuals’ FCs and SCs (out of 420 total) for 20,000 epochs with batch size *n* = 10 and learning rate *α* = 1*e −* 5. The final model was tested on the remaining 80 individuals, see Supplemental Figure 1 for a violin plot showing the correlation between predicted and true FC and correlations between inter-subject predicted FC, which were similar to the original publication’s results (20). Once trained and validated on the NeMo Tool’s controls, the neural network was then applied to the eSCs from each of the individual lesion masks to produce an eFC matrix for each pwMS. Regional eFC node strengths were calculated by taking the sum of each of the columns in the eFC after removing the negative entries.

### 2.3 Mass univariate analysis

First, demographics and clinical variables were tested for differences between the no disability vs evidence of disability groups using the Chi-squared test for qualitative variables and Wilcoxon rank-sum test for quantitative variables. Second, the pairwise and regional SC, FC, eSC, and eFC were tested for differences between the two disability groups using the Wilcoxon rank-sum test. Differences were considered significant when p *<* 0.05 after Benjamini-Hochberg (BH) correction for multiple comparisons (35). Entries in the eSC and SC were only considered in the univariate tests if they were non-zero for a majority of pwMS. All statistical analyses were performed and graphs created using R version 3.4.4 and Matlab version R2020a.

### 2.4 Classification analysis

Logistic regression with ridge regularization was used to classify pwMS into disability categories based on demographics/clinical information (sex, age, race, disease duration, clinical phenotype, and spinal lesion burden) and one of the pairwise or regional connectivity metrics derived from advanced MRI (SC and FC) or estimated from clinical imaging and the NeMo Tool (eSC and eFC).

The models were trained with outer and inner loops of k-fold cross-validation (k = 5) to optimize the hyperparameters and test model performance. The folds for both inner and outer loops were stratified to ensure that each fold contained the same proportion of subjects in the two classes as the original dataset. The inner loop (repeated over 5 different partitions of the training dataset only) optimized the set of hyperparameters that maximized validation AUC. A model was then fitted using the entire training dataset and those optimal hyperparameters, which was then assessed on the hold-out test set from the outer loop. The outer loop was repeated for 100 different random partitions of the data (See Figure 4 in the supplementary document). The median of AUC (overall 5 folds x 100 iterations = 500 test sets) was calculated to assess the performance of the models. In addition to the AUC values; Brier scores, sensitivity, specificity, and balanced accuracy are provided. The classification performances of different models were compared using permutation test (36) of 1000 permutations. P-values were the number of permutations that had a difference in means bigger than the original difference and were considered significant when p *<* 0.05 after BH correction for multiple comparisons (35). The important variables were identified using both univariate statistics (t-test or Wilcoxon rank sum test based on the connectivity type) and the mean feature weights (the beta parameter coefficients) over all 500 models (100 partitions of the data into 5 folds) (37). Regional and region-pair feature weights were also summarized at a functional network level by assigning each of the 68 cortical regions to one of seven canonical functional networks (38); subcortex and cerebellum were also added as networks. Pearson’s correlation coefficient was used to assess similarity of the feature weights from the four estimated and observed connectivity model pairs. Finally, as a sensitivity analysis, we repeated the classification analysis with EDSS of greater than 3 as the threshold for defining higher disability.

When the data contains class imbalance, models tend to favor the majority class. Due to the class imbalance in our data (67 vs 33 pwMS with no disability vs evidence of disability), the over-sampling approach Synthetic Majority Over-sampling Technique (SMOTE) (39) was used to obtain a balanced training dataset during cross-validation. SMOTE compensates for imbalanced classes by creating synthetic examples using nearest neighbor information and has been shown to be among the most robust and accurate methods with which to control for imbalanced data (40).

**Data/code availability statement:**

The deidentified data that support the findings of this study are available upon reasonable request from the corresponding author. The codes that were used in this study are publicly available. Please see https://github.com/cerent/MS-Estimated-Observed-SC-FC for the classification and statistical analyses as well as the violin plots, https://github.com/zijin-gu/sc2fc for the codes of the deep neural network framework used to estimate FC from eSC, and https://github.com/kjamison/connviewer for the circle and glass brain plots.

**Ethics statement:**

All studies were approved by an ethical standards committee on human experimentation, and written informed consent was obtained from all patients.

## 3 Results

### 3.1 Patient Characteristics

The 67 pwMS who did not have disability were significantly younger than those 33 pwMS having disability (corrected *p <* 0.05), and had a trend for shorter disease duration (corrected *p <* 0.10). The pwMS with progressive diagnoses were in the disability group, while those with CIS were in the no disability group. The population average lesion mask showed that lesions tended to occur most commonly in periventricular white matter, see Supplementary Figure 2.

**Table 1:** Subject demographics and clinical information. Values were presented as median [1st, 3rd quantile] for the continuous variables as the number/percent for sex. The two sets of groups (no disability vs evidence of disability) were tested for differences; p-values shown were corrected for multiple comparisons; *indicates significance at *p <* 0.05.

|  | All pwMS<br>(n=100) | No disability<br>(n=67) | Evidence of disability<br>(n=33) | p-value |
| --- | --- | --- | --- | --- |
| Age | 46 [37, 56] | 40 [35, 50] | 56 [46, 58] | 0.0001* |
| Female (%) | 66 (66%) | 46 (69%) | 20 (61%) | 0.56 |
| Disease Duration | 11 [7,16] | 10 [7,15] | 13 [9,17] | 0.06 |
| EDSS | 1 [0, 2] | 0 [0, 1] | 2 [2, 3] | <2.2e-16* |
| Number of spinal cord lesions | 1 [0,3] | 1 [0,3] | 2 [0,3] | 0.46 |
| Disease Category | 7 CIS, 88 RRMS,<br>5 Progressive MS | 7 CIS, 81 RRMS | 7 RRMS, 5 Progressive<br>MS | < 2.2e-16* |
| Lesion Volume (mm<sup>3</sup>) | 2065 [717, 4779] | 1995 [734,4200] | 2482 [453, 7788] | 0.49 |

### 3.2 Comparison of connectivity between disability groups

There was no significant difference in the pairwise or regional SC, FC, eSC, or between the regional eFC between the no disability vs evidence of disability groups. However, the pairwise eFC between the right rostral anterior cingulate and right temporal pole was significantly greater in pwMS who had no disability compared to those with evidence of disability (corrected p-value=0.04). Figures 2 and 3 illustrate the p-values (before correction) of group differences in pairwise and regional connectivity metrics between two disability groups, respectively. FC between ventral attention and frontoparietal and between cerebellum and subcortex, and eFC between ventral attention and visual were the connections that showed the biggest difference between pwMS who had evidence of disability vs those who had no disability. Regional node strength differences are shown in Figure 3; weaker eSC in all networks (except the cerebellum) and stronger eFC in all networks was observed in those pwMS who had evidence of disability. Observed FC was stronger in most networks (except subcortex, cerebellum and ventral attention) while observed SC was weaker in cerebellum and subcortex and stronger in limbic networks in pwMS who had evidence of disability.

**Figure 1:**
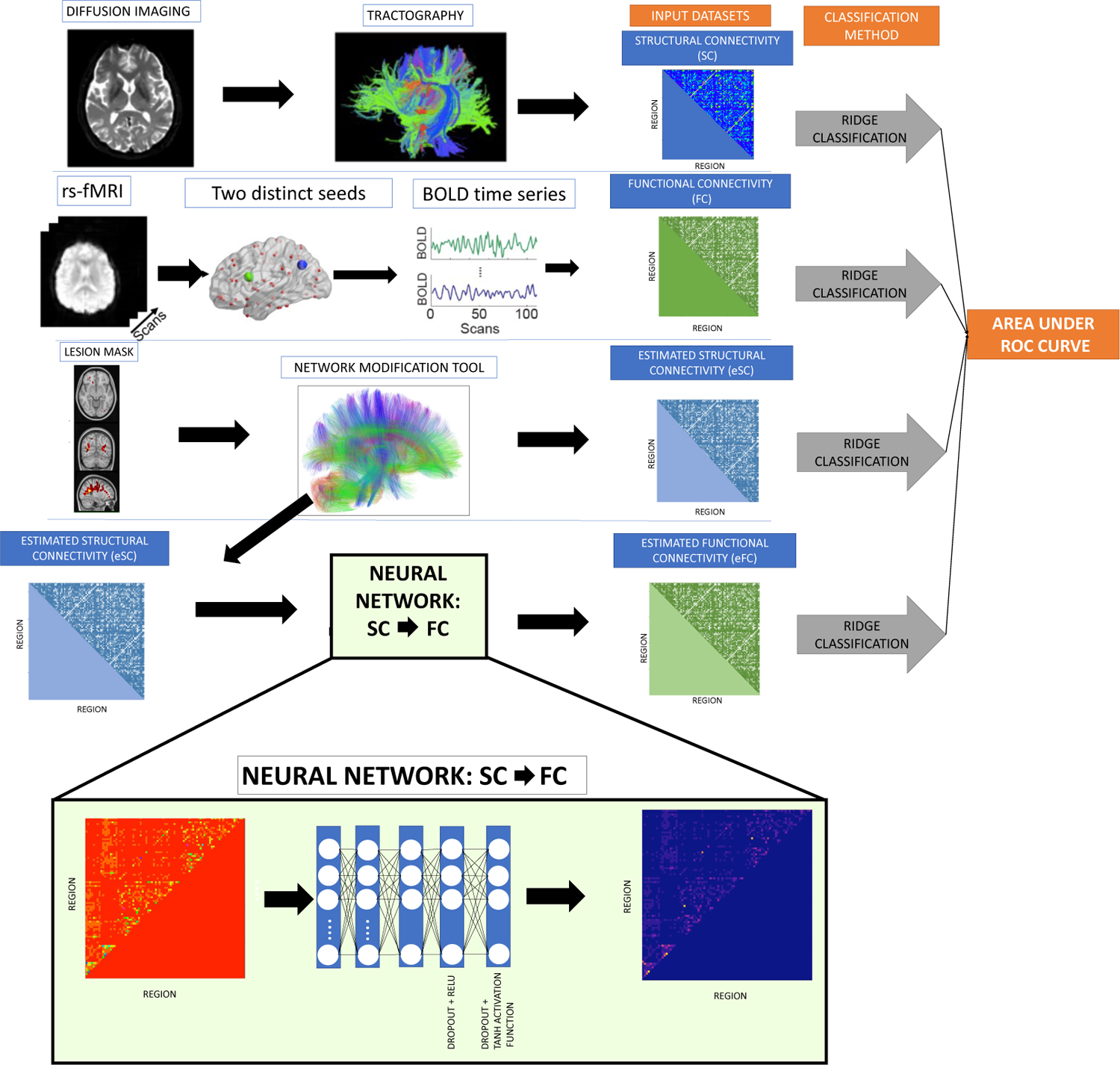
Workflow of the study. (A) Observed SC and FC were extracted from dMRI and fMRI directly in pwMS. The eSC and eFC matrices were computed using the lesion masks extracted from T2 FLAIR images and the Network Modification (NeMo) Tool. The eSC matrix was estimated by overlaying the lesion mask on a reference set of tractography results and removing streamlines that passed through the lesion mask. A deep neural network trained to predict FC from SC using the NeMo Tool’s control data (see tan box) was used to predict eFC rom eSC.

**Figure 2:**
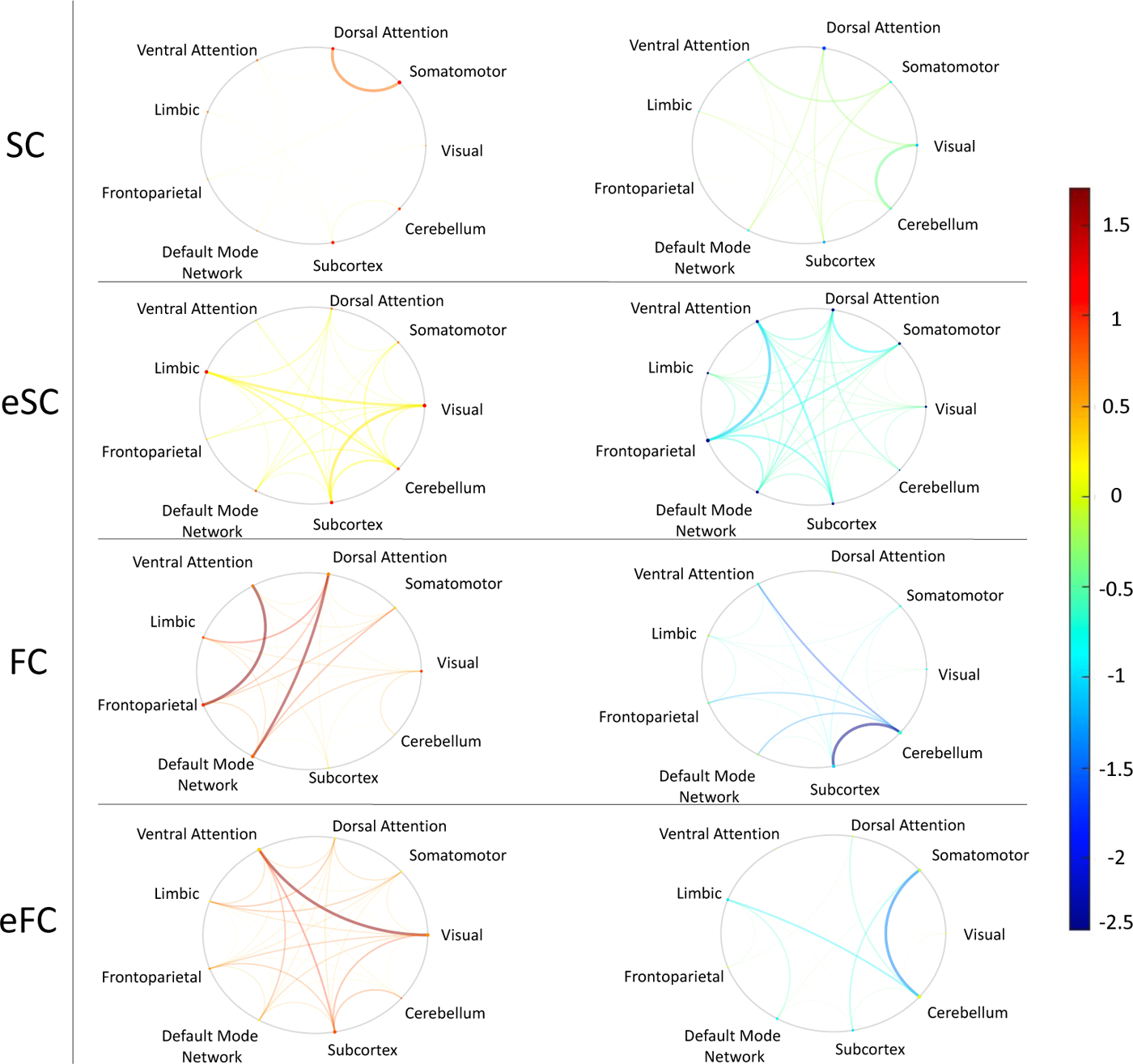
Comparison of pairwise connections between disability groups. The difference in connections between the evidence of disability vs no disability groups, where positive values (hotter colors) indicate pwMS who had evidence of disability had higher connectivity than those without disability, while negative values (cooler colors) indicate pwMS who had evidence of disability had weaker connectivity than those without disability. Group differences are visualized via -log(p)*sign(difference in mean or median), where p here is the uncorrected p. The p-values were obtained using t-test to compare FC and eFC between disability groups, while Wilcoxon rank-sum test was performed to compare SC and eSC values between disability groups. Difference in mean was used for FC and eFC, while difference in median was computed for SC and eSC. *Note: none of the pairwise connections for SC, FC or eSC were significant; only one eFC connection (between right rostral anterior cingulate and right temporal pole) was significantly larger in pwMS who had no disability.

**Figure 3:**
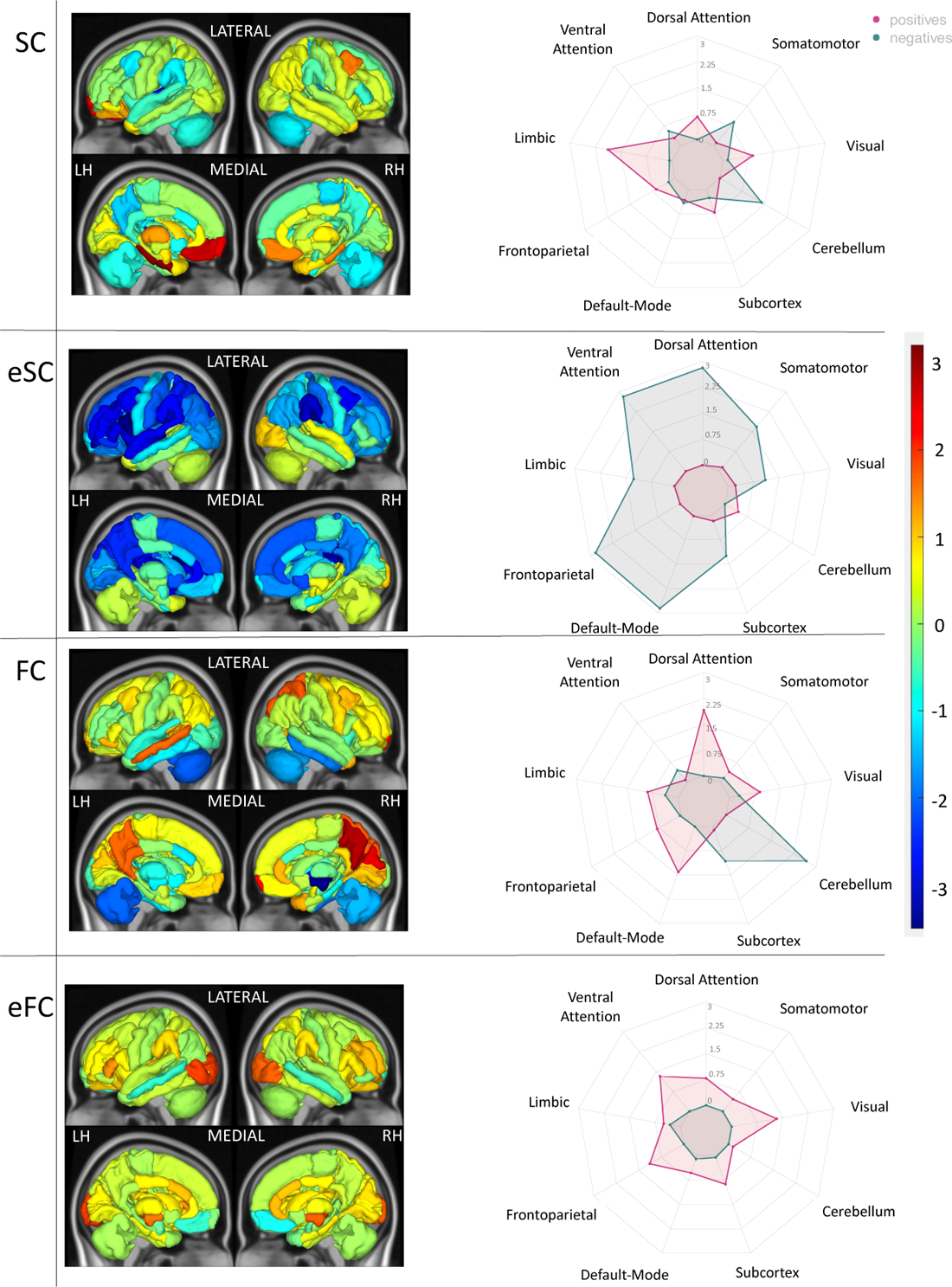
Comparison of node strength between disability groups. The difference in node strength between the evidence of disability vs no disability groups, where positive values (hotter colors) indicate pwMS who had evidence of disability had higher connectivity than those without disability, while negative values (cooler colors) indicate pwMS who had evidence of disability had weaker connectivity than those without disability. The colors display -log(p-value)*sign(difference in group means), where the p-value is uncorrected significance of the group difference via an unpaired t-test. The radial plots summarize network-level differences via the mean of the positive and negative -log(p)*sign(differences in group means) across 7 networks from the Yeo atlas, plus cerebellum and subcortex. The absolute value of the average negative connections were presented in the radial plots.

### 3.3 Classification results

Figure 4 shows that models based on pairwise SC and eSC had similar AUC (p=0.21, BH corrected), while pairwise eFC significantly outperformed FC (p-value*<*10e-4) and had the highest AUC of all the pairwise models. Both models based on estimated regional node strength outperformed observed regional node strength in classifying pwMS into disability groups (corrected p-value*<*0.01 for both regional SC vs eSC and regional FC vs eFC). The highest AUC over all 8 models tested was the one based on regional eFC, with a median of 0.681. Regional eSC and eFC had higher AUC results than pairwise eSC and eFC, and, while pairwise eFC outperformed pairwise eSC (corrected p-value*<*10e-4), regional eSC and eFC were not significantly different (corrected p-value=0.17). Sensitivity, specificity, balanced accuracy, and Brier score largely agreed with AUC results (see Supplementary Figure 3). Classification models were also tested using EDSS of 3 as the threshold for defining higher disability; the results are largely similar but the models had overall slightly larger AUCs (see Supplementary Figure 5).

**Figure 4:**
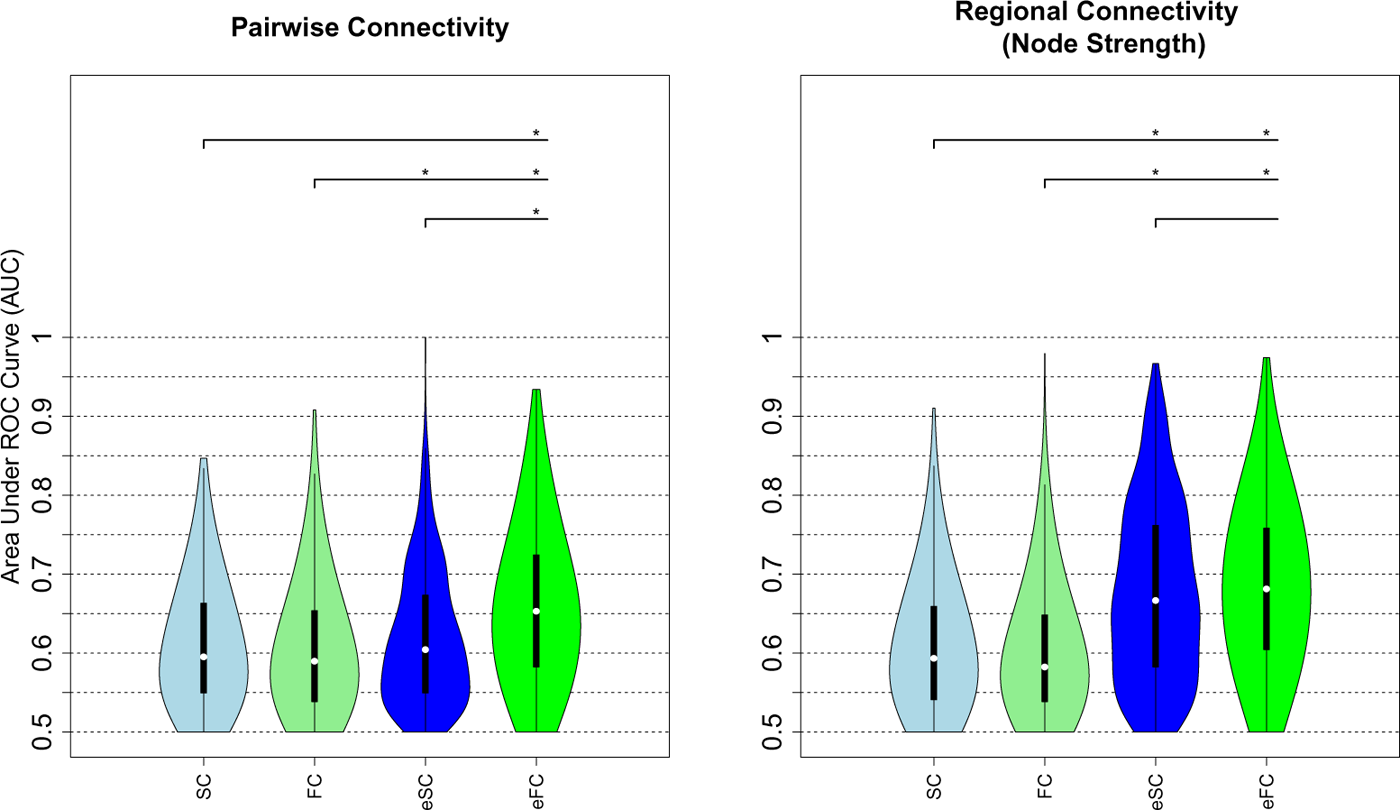
Classification results. AUC results obtained using pairwise connectivity (left panel) and regional node strength (right panel) to classify pwMS into no disability vs evidence of disability groups. The plots show the median (white dot) and inter quantile range (black bar) of the AUCs over the 500 hold-out test sets. The asterisks show the models that were significantly different after BH correction. SC: structural connectivity, FC: functional connectivity, eSC: estimated structural connectivity, and eFC: estimated functional connectivity

### 3.4 Feature weights

Age and sex had the highest weights in the pairwise and regional eSC and eFC models; being older or male was associated with having evidence of disability. Figure 5 shows the feature weights for the pairwise and regional eSC and eFC classification models. Connections from the cerebellum to the somatomotor regions and from cerebellum to limbic regions had the highest weights in both pairwise eSC and eFC models but in opposite directions; greater eFC and weaker eSC were associated with no disability. Additionally, weaker eSC between ventral attention and frontoparietal and subcortical networks was associated with evidence of disability while weaker eFC between cerebellum and somatomotor/visual/dorsal attention networks was associated with evidence of disability.

**Figure 5:**
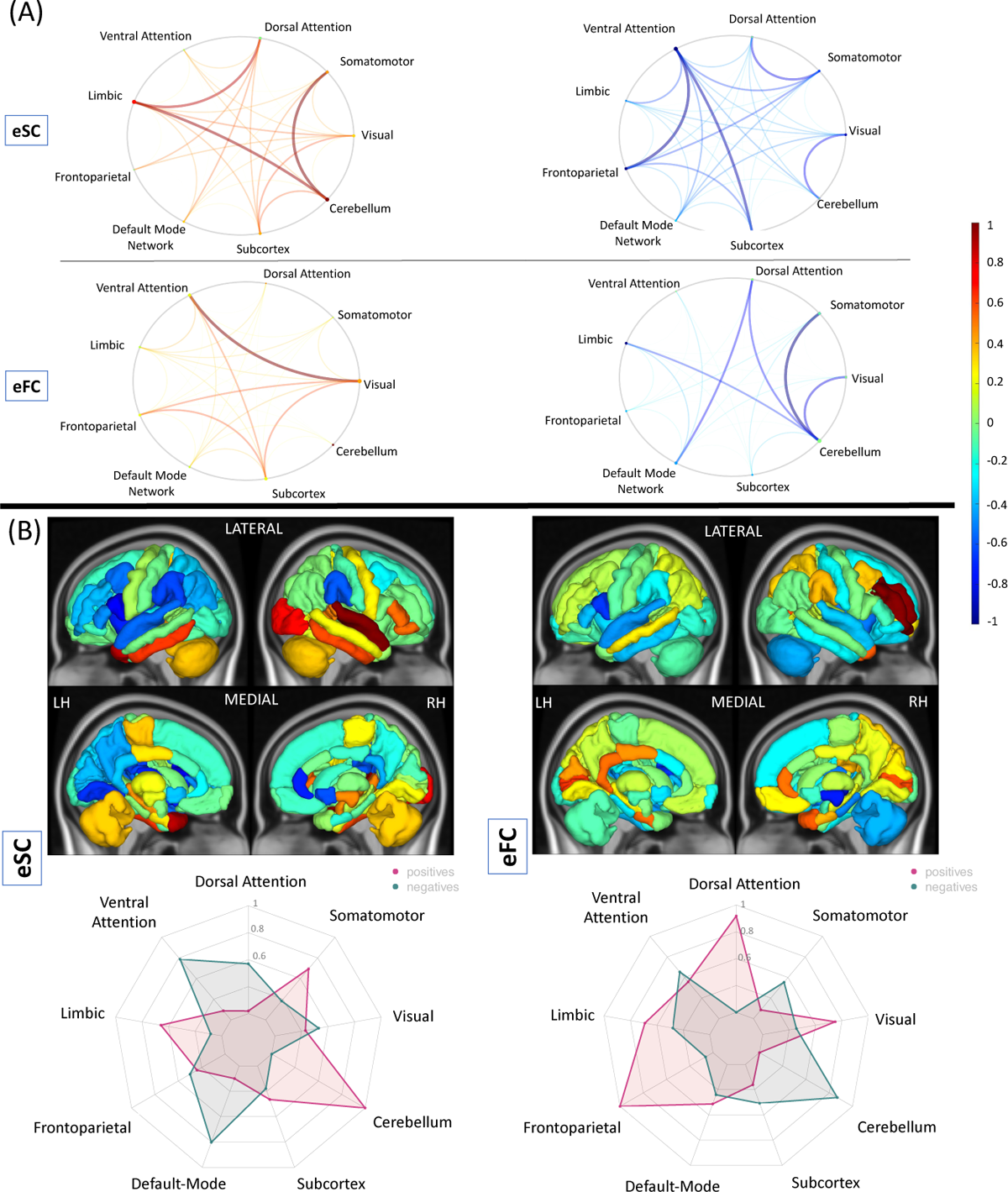
The feature weights computed with eSC and eFC models. The relative weights of the (A) pairwise and (B) regional (node strength) eSC and eFC obtained in classifying pwMS according to disability. Feature weight metrics were divided by the maximum of the absolute value over all models. Positive weights (hotter colors) indicate those connections/regions where stronger connectivity was associated with having evidence of disability class, while negative weights (cooler colors) indicate those connections/regions where stronger connectivity was associated with being in the no disability class. The circle plots summarize the pairwise feature weights by representing the mean of the positive and negative pairwise feature weights between functional network assignments (7 networks from the Yeo atlas, plus cerebellum and subcortex). The radial plots summarize the feature weights of regional models by representing the mean of the positive and negative weights in the same functional networks used for the circle plot. The absolute value of the average negative connections were presented in the radial plots.

Weaker eSC node strength in the cerebellum, somatomotor, and limbic networks and stronger eSC node strength in the ventral/dorsal attention and default mode networks were associated with no disability. Stronger eFC node strength in the visual, dorsal attention and frontoparietal was associated with evidence of disability, while weaker eFC node strength in the cerebellum and somatomotor networks was associated with evidence of disability.

Figure 6 shows the feature weights for the observed SC and FC models, which both had lower classification accuracy and only provided them here for comparison to the eFC/eSC models. Weaker SC between the dorsal attention and visual networks and weaker FC between cerebellum and subcortical, ventral attention and default mode networks were associated with greater disability. Weaker SC between dorsal attention and cerebellum/somatomotor networks and weaker FC between default mode and dorsal attention networks was associated with no disability. Weaker SC and FC to/from the right cerebellum to the rest of the brain was also found to be associated with having evidence of disability (41). There was a moderate correlation between the feature weights from the regional FC and eFC models (r=0.52), while the correlations between pairwise FC vs eFC, as well as pairwise/regional SC vs eSC were non-significant and weak (r*<*0.1) (See Supplementary Table 1). When the features were analyzed at network level, weaker SC and FC in the cerebellum was commonly found as associated with disability.

**Figure 6:**
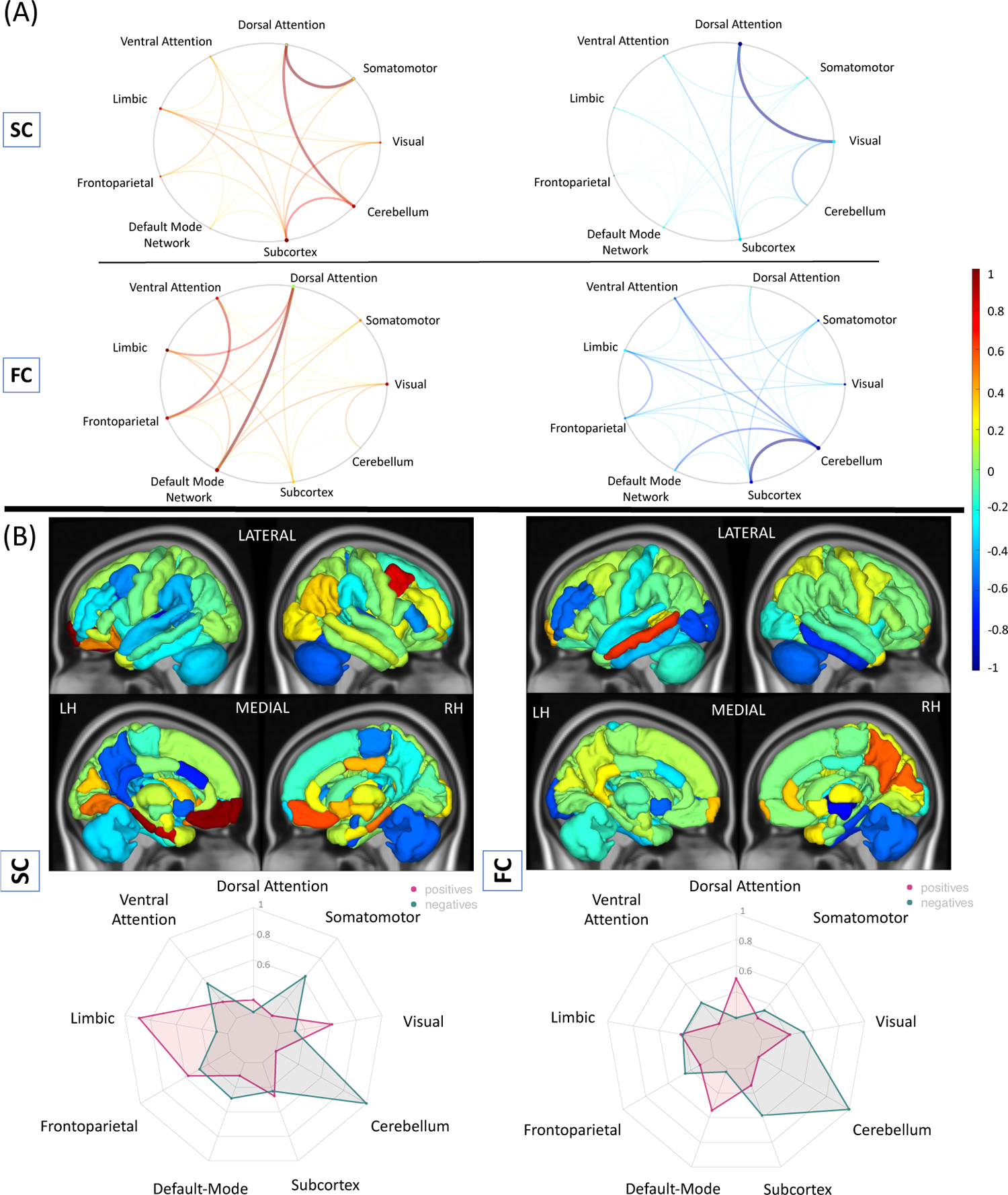
The feature weights computed with SC and FC models. The relative weights of the (A) pairwise and (B) regional (node strength) SC and FC obtained in classifying pwMS according disability. Feature weight metrics were scaled by the maximum of the absolute value over all models. Positive weights (hotter colors) indicate those connections/regions where stronger connectivity was associated with having evidence of disability class, while negative weights (cooler colors) indicate those connections/regions where stronger connectivity was associated with being in the no disability class. The circle plots summarize the pairwise feature weights by representing the mean of the positive and negative pairwise feature weights between functional network assignments (7 networks from the Yeo atlas, plus cerebellum and subcortex) The radial plots summarize the feature weights of regional models by representing the mean of the positive and negative weights in the same functional networks used for the circle plot. The absolute value of the average negative connections were presented in the radial plots.

## 4 Discussion

In this study, we compared the performance of models using observed SC and FC (derived from advanced MRI directly in pwMS) against models using estimated SC and FC (derived from clinically-acquired lesion masks and the NeMo Tool) in classifying pwMS into no disability vs evidence of disability groups. We also identified the regions and/or connections that were most different between and important in classifying pwMS into disability groups and compared feature weights over the estimated and observed models. Our main findings were (1) models based on regional eSC and pairwise/regional eFC outperformed models based on their observed counterparts, (2) models based on pairwise eSC performed just as well as models based on pairwise observed SC, (3) models based on regional eSC and eFC performed the best out of all models considered (4) stronger eFC node strength of regions in the visual network and weaker eSC node strength of regions in the default mode and ventral attention networks was associated with disability, and (5) feature weights computed from the regional eFC and FC models were moderately, significantly correlated.

### Previous studies comparing estimated and observed connectivity

The original NeMo Tool was used previously to map disruptions of SC to concurrent and future impairments in stroke and MS (3; 2; 42; 43). As far as we know, no study to date has compared indirectly estimated SC or FC from the NeMo tool or any other techniques to directly observed SC or FC in their respective abilities to predict impairment or disability in pwMS. However, we are aware of several recent studies in stroke that have shown contrasting results. For example, one study showed that lesion masks themselves, in addition to estimated structural disconnectivity and to a lesser extend directly observed FC, predicted multiple post-stroke impairments while estimated functional network disruptions did not (23). In a follow-up replication study with the same dataset, Cohen et al. (44) found that modifying the way in which estimated functional disruption was calculated resulted in models with explained variance similar to the estimated structural disconnectivity. Although our estimated FC was calculated in a very different manner than these studies, our results agree with the latter one. In fact, our results in pwMS actually show that models based on pairwise eFC outperform models based on pairwise eSC. Our novel approach to estimating FC from lesion masks could be a complimentary or alternative technique to lesion-network dysfunction mapping that may allow even greater explained variance when performing brain-behavior mapping in individuals with neurological disease.

### Estimated vs observed connectivity: advantages and disadvantages

Collecting and processing dMRI and fMRI in patient populations is difficult, as individuals often cannot tolerate long scan times, scans are expensive and processing of the images requires a high level of expertise. In addition, anatomical or physiological changes like edema and/or inflammation can add noise to already noisy dMRI and fMRI modalities and further corrupt their accurate measurement. Finally, big data approaches, including deep learning, require very large samples of individuals in order to create models that can generalize to new patient populations and truly be informative of disease mechanisms or be used for accurate prognoses. Collecting advanced MRI in the number of people needed to train these types of models would be nearly impossible, but our approach to estimating both SC and FC from clinically-acquired lesion masks is much more feasible. Additionally when obtaining large sample sizes, data is often combined across different sites. Inter-site variability in advanced MRI acquisitions is much more prominent than would be anatomical imaging used to create lesion masks only. Therefore, the NeMo tool can be advantageous as it is easier to implement, provides less noisy data, and allows to study larger number of patients compared to advanced imaging techniques, because only clinically-acquired lesion masks are used to estimate SC and FC networks.

The NeMo Tool does have some disadvantages compared to collecting advanced MRI directly in patient populations. First, the eSC and eFC are based on a population of 420 controls and may not closely reflect the particular individual’s anatomical or physiological connections. Second, there may be compensatory rewiring of SC or reorganization of FC that will not be reflected in the eSC and/or eFC, which means that use of the NeMo tool may not be appropriate in studies aimed at identifying recovery mechanisms. Third, there may be other pathologies in the brain that are not included in lesion masks and are thus not considered by the NeMo Tool. Finally, lesion masks must be coregistered to a common space which can be inaccurate at times in individuals with brain atrophy or gross abnormalities (45).

### Comparison to previous classification studies in pwMS

Previous studies have used SC and FC metrics to distinguish pwMS from HC and/or to classify pwMS by disability level defined by thresholding EDSS scores (46; 47; 48). A previous study compared the classification accuracy of SC and FC to discriminate pwMS into disability groups using a threshold EDSS of 2 (47); our models based on observed connectivity metrics had higher AUC (0.47 vs 0.59 for models based on SC and 0.53 vs 0.59 for models based on FC). The difference in results might be due to different clinical stage used in the studies (they used relapsing remitting and our study that included all phenotypes), different classification methods (they used support vector machines and we used a ridge classifier) and different MRI acquisition and processing protocols. In a recent preprint, we used the NeMo Tool’s estimated regional disconnectivity measures based on different lesion types (hyperintense rim positive or negative on quantitative susceptibility mapping imaging) and found AUCs similar to those found here (0.63-0.67) in classifying a different population of pwMS into no disability vs evidence of disability categories also based on a threshold EDSS of 2 (41). An EDSS threshold of 3 was identified as the cut-off for functional reorganization and adaptation in MS (14). This study found no significant difference in FC between HC and pwMS who had an EDSS *≤* 3, while FC decreased in the pwMS who had an EDSS *>* 3. Our replication study using a threshold EDSS of 3 and found largely similar results (see Supplementary Figure 5).

### Comparison of feature weights: estimated vs observed connectivity models

Even though the correlation between the raw values of regional FC and eFC was weak (r=0.09, p-value *<* 10e-16), the feature weights from the regional FC and eFC models are moderately correlated (Pearson’s r = 0.52, p-value *<* 10e-7). In particular, increased FC and eFC in cerebellum, subcortex, somatomotor, and ventral attention networks were commonly found to be associated with no disability while decreased FC and eFC in dorsal attention and default mode networks were commonly found to be associated with no disability. Moreover, higher node strength in many regions (*>*35) in the FC and eFC models were commonly associated with disability. This finding is consistent with previous studies showing upregulation of FC reflecting possible compensation in pwMS having lower EDSS compared to those with greater EDSS (14).

### Assessment of feature importance

A recent study (37) showed that coefficients of models predicting cognition and sex using observed FC can be unreliable. In their paper, they argue that increasing sample size, applying the Haufe transformation and using non-sparse regularization can all improve the feature weight reliability; furthermore they found that mass univariate results can be more reliable than model coefficients. In our study, we used (non-sparse) ridge regularization, however the Haufe transformation cannot be applied as our model contains binary/categorical demographic variables (such as sex and spinal cord lesion category). Therefore, here we focus on reporting classification model coefficients that also agreed with the mass univariate group comparison results.

### eSC in the default mode and ventral attention and eFC in the visual networks had the highest importance for disability classification in MS

In our study, weaker eSC in the default mode and ventral attention networks was commonly found as associated with disability in both univariate test and the classification model. Structural disruption and functional alterations in the default mode network were previously found as associated with poorer cognitive abilities and cognitive rehabilitation outcomes (49; 22). However, this is the first study that showed the relationship between disability and connectivity changes in these networks in MS. In our study, we also showed that greater eFC in the regions of the visual network was associated with disability. Our results were in concordance with a previous study (50) that also showed greater FC of the visual network in pwMS as compared to HCs and interpreted as the presence of cross-modal plasticity mechanisms after visual impairment seen in early MS (51).

### eSC and eFC between the cerebellum and somatomotor networks is central to accurate disability classification

It was not surprising that the eSC and eFC from the somatomotor network and cerebellum had the highest weights in identifying pwMS’s disability level, as these networks are central to motor and movement. In addition to the cerebellum-somatomotor/limbic connection, a decrease in the connectivity metrics in the cerebellum was also associated with having more probability of having disability. Previous studies have shown relationships between cerebellar pathology and impairments in motor control and cognition (52; 53). The presence of cerebellum-related symptoms at the onset of MS such as coordination issues or tremor were i) shown to be associated with shorter time to an EDSS of 6 (54) and ii) related to earlier onset of progressive disease diagnosis (55). Atrophy in the anterior cerebellum was associated with motor dysfunction (52) in pwMS. In our previous study, the cerebellum was also shown as the most important region in classifying pwMS by disability level, where eSC was computed using only hyperintense rim lesions from quantitative susceptibility imaging (41). Another study that investigated FC in the cerebellum found that the loss of functional cerebellar connections was related to disability. Our results and previous findings may suggest that decreasing structural deterioration and functional upregulation in the cerebellum may be a new target for clinical trials (14).

Among demographics and clinical variables, age and sex appeared to be an important predictor for disability classification; specifically, being older and male was associated with a higher probability of being in the disability group. It has been shown previously that male patients tend to have more severe disease onset with accelerated clinical progression in MS (56).

### Limitations and Future Work

One of the limitations of the present study was in the quality of the advanced MRI collected directly in pwMS. The fMRI acquisition time was relatively short and had a longer TR (6 min, TR = 2.3 s) and the dMRI acquisition had only a single b-value of 800 and only 55 directions. These limitations may mean less accurate observed FC or SC, which could have negatively impacted their classification accuracy. A future study will be needed to compare eSC and eFC to SC and FC from higher quality fMRI and dMRI acquisitions, however if high-quality SC and FC are superior than eSC and sFC, it may still be preferable to use estimates given feasibility issues. The estimates of SC and FC from the NeMo tool are based on a database of controls and thus cannot capture an individual’s specific SC anatomy or the particulars of the relationship between SC and FC under the influence of lesion pathology. Additionally, there may be pathologies in the white matter of pwMS that are not contained in the lesion masks and are thus not considered in the NeMo Tool’s estimates. However, MS lesions disrupt diffusion MRI signals and add noise to tractography results, and the current findings indicate that the NeMo Tool is a good alternative to performing tractography directly in pwMS. This study investigates cross-sectional relationships between disability and connectivity; a future study mapping baseline connectivity to future disability is be needed to better understand the prognostic ability of estimated connectomes.

## Conclusions

This is the first study to show that models using estimated SC and FC based only on lesion masks and the NeMo Tool outperform models based on observed SC and FC extracted from advanced MRI in classifying pwMS into disability categories. Models based on eFC’s regional node strength had the highest overall classification performance. Stronger eFC node strength of regions in the visual network and weaker eSC node strength of regions in the default mode and ventral attention networks was associated with disability. A deeper understanding of the role of the connectome in MS is needed if we are to gain a comprehensive view of the disease, develop more accurate prognostic or therapeutic tools and ultimately improve clinical outcomes in pwMS. This work provides a viable alternative to performing high-cost, advanced MRI in patient populations, brings the connectome one step closer to the clinic.

## Abbreviations

AUC: area under curve

EDSS: extended disability status score

eFC: estimated functional connectivity

eSC: estimated structural connectivity

FC: functional connectivity

GM: gray matter

HC: healthy control

QSM: quantitative susceptibility imaging

LST: lesion segmentation tool

MS: multiple sclerosis

NeMo: network modification

pwMS: people with multiple sclerosis

SC: structural connectivity

WM: white matter

## Acknowledgements

C.T. helped with image post-processing, carried out the statistical analyses and wrote the article.

K.J. collected the data, performed pre- and post-processing of MRI data and reviewed the article.

Z.G. created the neural network to predict FC from SC.

S.G. collected the data, helped interpret results and reviewed the article.

A.K. designed and supervised the study, collected the data and edited the article.

## Citation gender diversity statement

We used classification of gender based on the first names of the first and last authors (Dworkin et al., 2020), with possible combinations including male/male, male/female, female/male, and female/female. The gender balance of papers cited within this work was quantified using gender-api.com. The authors with a gender estimation accuracy lower than 90% were checked using manual gender determination from authors’ publicly available pronouns. Among the 56 cited works, 2 articles had only one author. Among the 55 cited works with more than one author, 45% (n = 25) were MM, 35%(n = 19) were WM, 11% (n = 6) were MW, and 9% (n = 5) were WW.

