## Supplementary Material for "Estimated connectivity networks outperform observed connectivity networks when classifying people with multiple sclerosis into disability groups"

### 1 Supplementary Document

#### 1.1 Network Modification Tool

The Network Modification 2.0 (NeMo 2.0) tool estimates structural connectivity disruption due to a lesion or injury from a database of HC's structural connectivity. We first computed a database of whole-brain tractograms for 420 unrelated HC (206 female, 214 male, 28.7 ± 3.7 years) from the Human Connectome Project Young Adult (HCP-YA) dataset. The HCP diffusion data has 1.25mm isotropic voxels, 3 shells (b=1000,2000,3000) and 90 directions per shell, and was collected with both R-L and L-R phase encoding. HCP data have been minimally preprocessed to correct for motion, EPI and eddy-current distortion, and registered to subject T1 anatomy (1). We used MRtrix3 to estimate a voxel-wise multi-shell, multi-tissue constrained spherical deconvolution (CSD) model (2), followed by whole-brain deterministic (sd-stream) tractography with MRtrix3 (3) with dynamic white-matter seeding (4). Streamlines for each HCP subject were warped into a common volumetric space (MNI152) to create the final reference set.

Given a lesion mask in MNI152 space, the NeMo tool identifies all streamlines that pass through the lesioned voxels and removes them before calculating the eSC matrix by taking the sum of the SIFT2 weights between remaining streamlines connecting each pair of regions in the pre-selected atlas and dividing by the sum of the region-pair volumes. The NeMo Tool also computes an estimated pairwise and regional structural disconnectivity score for each gray matter region-pair or region in a given atlas, representing the fraction of streamlines connecting those pairs of regions or connecting to that region that pass through the lesion. The eSC matrix and disconnectivity measures are calculated for each of the 420 HCP database subjects and then averaged across the subjects to create the final eSC and/or disconnectivity estimate for each lesion mask. In this paper, we use solely the eSC as to compare with the observed SC extracted from dMRI in the individual pwMS.

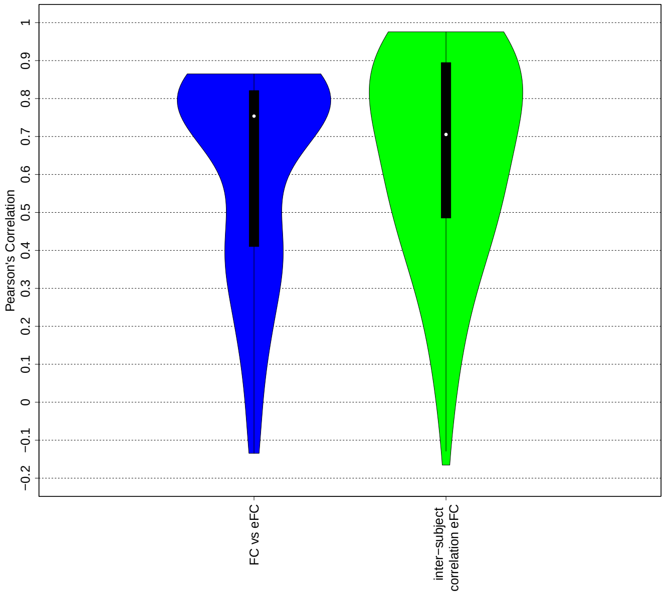


Figure 1: The violin plot of the Pearson's correlation metrics computed between FC and eFC and inter-subject correlation between eFCs obtained from deep neural network using HCP dataset.

#### 1.2 Average lesion mask

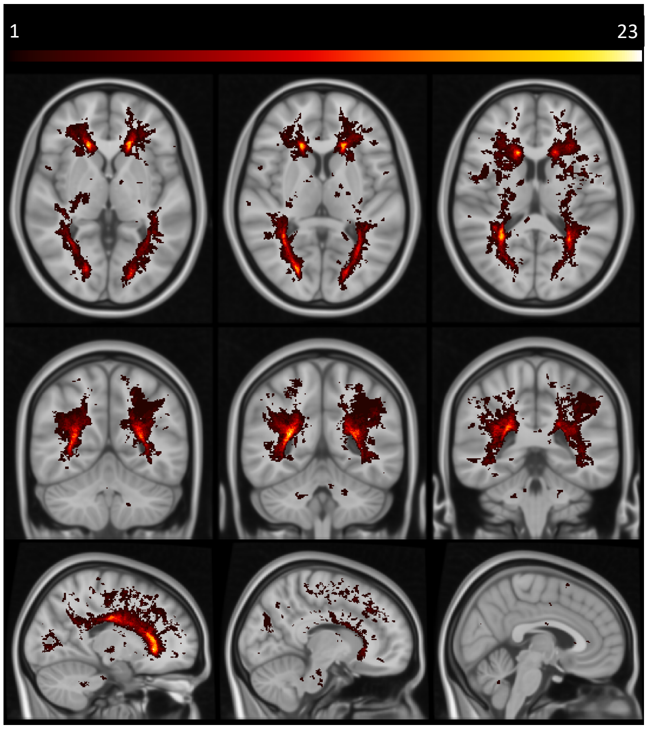


Figure 2: Average T2FLAIR lesion mask across 100 people with MS. Color indicates the number of individuals (out of 100) that had a lesion in that voxel.

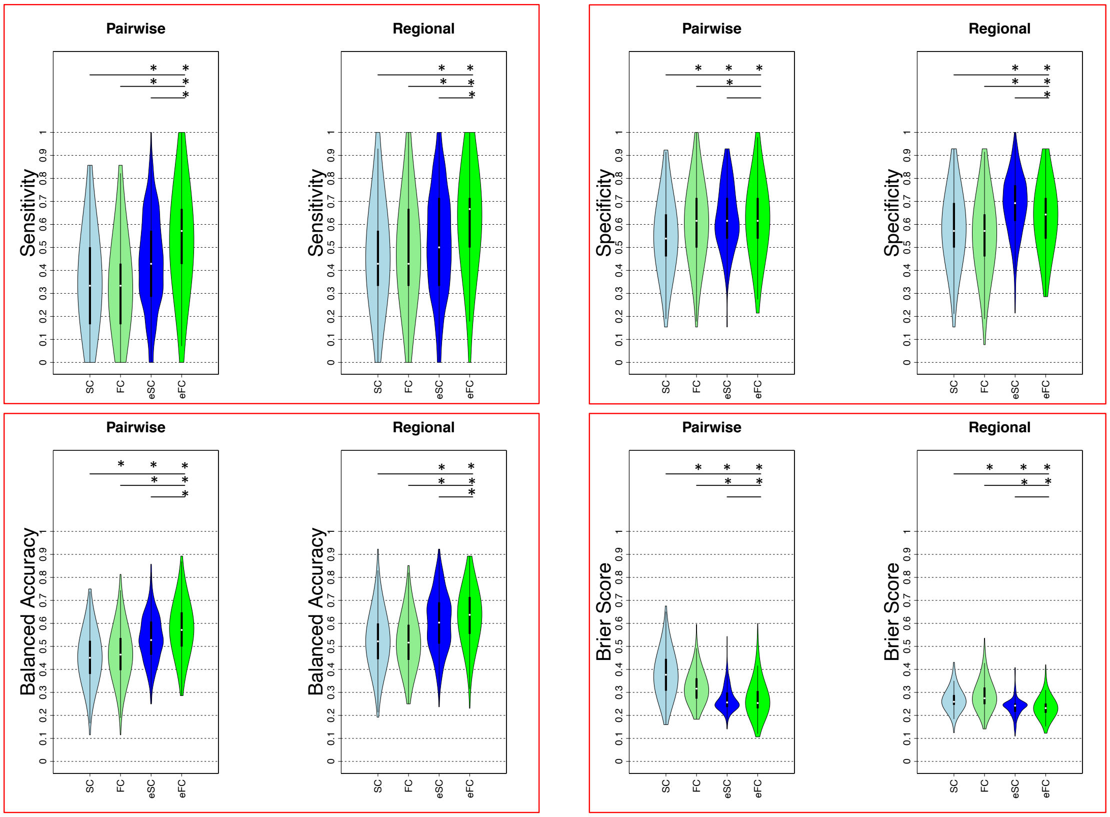


Figure 3: The sensitivity, specificity, balanced accuracy, and Brier score results. EDSS 2 was used as a threshold to define the no disability vs an evidence of disability groups.

#### 1.3 Classification analysis

##### 1.3.1 Logistic regression with ridge regularization

Linear regression is one of the most widely used statistical methods available today. However, over-fitting problem may occur when performing linear regression with high dimensional data. Ridge regression estimates the parameter coefficient ($\hat{\beta}_{ridge}$) by applying an $L_2$ penalty as follows:

$$\hat{\beta}_{ridge} = (X^T X + \lambda I)^{-1} X^T Y \tag{1}$$

where $X$ is the input and $Y$ is the output.

$\lambda \geq 0$ is a tuning parameter for the penalty, which is determined separately. Our $\lambda$ was chosen to be the value in $[10^{-3}, 10^{+3}]$ using steps of 10 that maximized classification accuracy (AUC) in the inner loop on the validation set. Once identified, this value was used to fit the model in the outer loop..

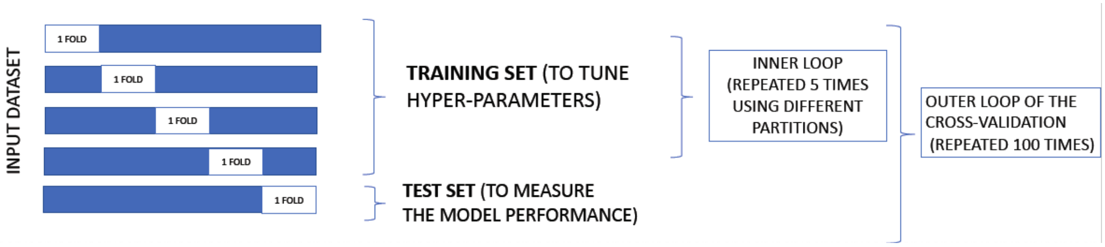


Figure 4: The nested cross-validation approach used in this study.

We considered feature weights in the ridge regression models to be the mean of the ridge regression coefficients ($\hat{\beta}_{ridge}$) from all 500 models (100 partitions of the data into 5 folds).

| | | **Pairwise** | **Regional** |
|---|---|---|---|
| **Raw values** | SC vs eSC | 0.81 (p-value<2.2e-16) | 0.62 (p-value<2.2e-16) |
| | FC vs eFC | 0.23 (p-value<2.2e-16) | 0.09 (p-value<2.2e-16) |
| **Feature weights** | SC vs eSC | 0.01 (p-value = 0.324) | 0.03 (p-value=0.729) |
| | FC vs eFC | -0.03 (p-value = 0.041) | 0.52 (p-value=2.569e-07) |

Table 1: The Pearson's correlation coefficients and p-values between raw pairwise/regional SC vs eSC and FC vs eFC values and correlations between feature weights from models based on those values.

##### 1.3.2 Classification results using a threshold EDSS of 3

Eighty-eight 88 pwMS had no disability and 12 pwMS had an evidence of disability when an EDSS of 3 was used as threshold for dividing the no disability vs an evidence of disability groups; Figure 5 shows the classification results. The AUC results were overall slightly higher than those obtained when using a threshold EDSS of 2. The pairwise eFC again outperformed other pairwise models with a median of 0.685, while regional eSC showed higher AUC results compared to regional eFC with a median AUC of 0.667 (corrected p-value=0.0276).

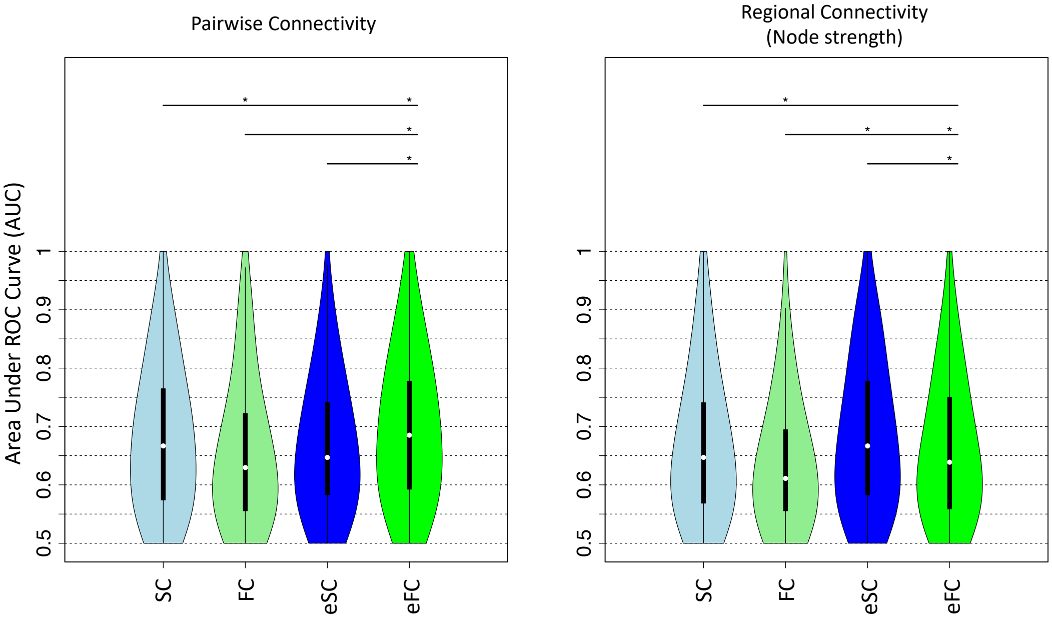


Figure 5: Classification results when using a threshold EDSS of 3 to define no disability vs an evidence of disability.

1
